# Allele-specific gene regulation by KDM6A

**DOI:** 10.1101/2020.09.09.289926

**Authors:** Wenxiu Ma, He Fang, Nicolas Pease, Galina N. Filippova, Christine M. Disteche, Joel B. Berletch

**Affiliations:** Department of Statistics, University of California Riverside, Riverside, California 92521, USA; Department of Pathology, School of Medicine, University of Washington, Seattle, Washington 98195, USA; Department of Genome Sciences, University of Washington, Seattle, Washington 98195, USA; Department of Medicine, School of Medicine, University of Washington, Seattle, Washington 98195, USA

**Keywords:** Epigenetics, histone methylation, imprinting, allele-specific, gene regulation, development, chromatin, parent-of-origin

## Abstract

KDM6A demethylates the repressive histone mark H3K27me3 and thus plays an important role in developmental gene regulation. KDM6A expression is female-biased due to escape from X inactivation, suggesting that this protein may play a role in sex differences. Here, we report that maternal and paternal alleles of a subset of mouse genes are differentially regulated by KDM6A. Knockouts of *Kdm6a* in male and female embryonic stem cells derived from F1 hybrid mice from reciprocal interspecific crosses resulted in preferential downregulation of maternal alleles of a number of genes implicated in development. Moreover, the majority of these genes exhibited a maternal allele expression bias, which was observed in both reciprocal crosses. Promoters of genes downregulated on maternal but not paternal alleles demonstrated a loss of chromatin accessibility, while the expected increase in H3K27me3 levels occurred only at promoters of genes downregulated on paternal but not maternal alleles. These results illustrate parent-of-origin mechanisms of gene regulation by KDM6A, consistent with histone demethylation-dependent and -independent activities.

## INTRODUCTION

Developmental processes are governed by specific patterns of epigenetic regulation of gene expression, which include enzymatic modifications of histone tails. For example, timely activation of *Homeobox (Hox)* genes via resolution of bivalency in early development is facilitated by the histone demethylase KDM6A that removes methylation at lysine 27 of histone H3 (Agger et al., 2007; De Kumar et al., 2015; Dhar et al., 2016; Yu et al., 2019). KDM6A has been implicated in many developmental pathways such as myogenesis, cardiac development, neural stem cell differentiation and immune cell functions (Bosselut, 2016; Itoh et al., 2019; Lee et al., 2012; Lei and Jiao, 2018; Seenundun et al., 2010). In addition to its demethylase activity, KDM6A can influence gene expression by associating with chromatin remodeling proteins and by regulating H3K27 acetylation levels at active enhancers through recruitment of p300 as part of the MLL complex (Miller et al., 2010; Shpargel et al., 2017; Wang et al., 2017). In addition to its major role in gene activation KDM6A can also act as a repressor of gene activity, highlighting the complexity of this protein (Gozdecka et al., 2018). The higher level of KDM6A in female mammals due to escape from X inactivation suggests a role in sex-specific gene regulation, which we previously demonstrated for the reproduction-related genes *Rhox6/9* (Berletch et al., 2013; Greenfield et al., 1998). KDM6A is encoded by a dosage-sensitive gene and has a Y-encoded homolog (UTY) that has little or no demethylase activity (Bellott et al., 2014; Cortez et al., 2014; Shpargel et al., 2012). Knockout of *Kdm6a* is homozygous lethal in female mice, while a small number of runted males survive, suggesting that UTY partially compensates for loss of KDM6A possibly via demethylation-independent mechanisms (Shpargel et al., 2014).

Here, we hypothesize that differential expression of KDM6A in male and female germline could potentially lead to parent-of-origin biased gene expression in the next generation. To address this possibility, we investigated the role of KDM6A in the epigenetic regulation of maternal and paternal alleles of genes in mouse embryonic stem (ES) cells. *Kdm6a* knockout (KO) was obtained by CRISPR/Cas9 approaches in ES cells derived from F1 hybrid mice from reciprocal crosses between C57BL/6 and *Mus castaneus* in which alleles can be distinguished based on SNPs. Genome-wide changes in allele-specific gene expression were measured by RNA-seq. Most of the genes with significantly reduced expression were differentially downregulated in an allele biased manner, with more genes downregulated on maternal compared to paternal alleles. Chromatin structure and epigenetic changes measured by allele-specific ATAC-seq and ChIP-seq for H3K27me3 suggested that, depending on the parental allele, gene regulation by KDM6A occurs through demethylase-dependent and/or demethylase-independent mechanisms.

## RESULTS

### KDM6A KO in ES cells results in expression changes in developmentally important genes

*Kdm6a* was edited using CRISPR/Cas9 in F1 male and female ES cells derived from a cross between C57BL/6J x *Mus castaneus* (BC) and in F1 male ES cells derived from the reciprocal *Mus castaneus* x C57BL/6J cross (CB), in which alleles can be distinguished using SNPs (Figure S1) (Barakat et al., 2011). As detailed in Methods we derived stable *Kdm6a*-targeted ES cell clones from both BC and CB crosses, using CRISPR-Cas9 to induce either a deletion between exons 2 and 4 (ΔE) or a deletion of exons 1 and 2 in the promoter region (ΔP) (Figure S1A-D). We verified *Kdm6a* editing and loss of KDM6A protein in eight KDM6A KO clones, including four male hemizygous clones (*Kdm6a*^*ΔE1*^, *Kdm6a*^*ΔE3*^, *Kdm6a*^*ΔP4*^, *Kdm6a*^*ΔP6*^, hereafter BC *Kdm6a*^*Δ/Y*^) from the BC cross, one of which (*Kdm6a*^*ΔP6*^, hereafter BC *Kdm6a*^*ΔP/-Y*^) has lost the Y chromosome, three male hemizygous clones (*Kdm6a*^*ΔE2*.*5*^, *Kdm6a*^*ΔE2*.*7*^, *Kdm6a*^*ΔP2*.*1*^, hereafter CB *Kdm6a*^*Δ/Y*^) from the CB cross, and one female homozygous clone (BC *Kdm6a*^*ΔP/ΔP*^) from the BC cross (Figure S1A-E; Table S1). An off-target 314 kb deletion of three adjacent genes (*Sh3kbp1, Map3k15, Pdha1*) was identified in the BC-derived *Kdm6a*^*ΔE1*^ and *Kdm6a*^*ΔE3*^ clones, consistent with a common origin of these lines (Figure S1F and data not shown). We also isolated control male ES cell clones (hereafter called CRISPR-controls) derived from the BC cross, which were subject to the CRISPR/Cas9 treatment but did not exhibit a deletion of *Kdm6a* (Table S1).

To identify genes targeted by KDM6A RNA-seq analysis was done to compare gene expression between wild-type (wt) and the KDM6A KO clones derived from the BC and CB crosses. Using principal component analysis and hierarchal clustering based on expression values for all transcribed genes, male clones segregated from female clones, and wt clones clustered separately from KO clones (Figure S2A, B). The lines derived from either BC or CB crosses clustered together, suggesting strain-specific differences. Note that the ES cells derived from the CB cross contain an actin-GFP transgene, which could also contribute to gene expression differences between the crosses. In addition, we cannot exclude clonal or batch effects. DESeq2 was employed to identify differentially expressed genes (DEGs) in each cross using an FDR cutoff of <0.05 and a fold change threshold ≥1.5 fold (Love et al., 2014).

In the BC cross we identified 195 downregulated and 334 upregulated DEGs in the three male KO clones that retain the Y chromosome (*Kdm6a*^*Δ/Y*^) compared to the two male wt clones (Figure 1A; Table S2). As expected, downregulated DEGs include known KDM6A targets (Figure S2C). Reduced expression of selected genes was confirmed by qPCR and RT-PCR (Figure S2D and S4B). Next, we compared our list of downregulated DEGs to a publicly available gene expression dataset obtained in independent KDM6A KO male mouse cell lines (Wang et al., 2017). After applying our expression fold change threshold and FDR cutoff to the published data, we found concordance for 79/195 (40%) and 104/334 (31%) of our downregulated and upregulated DEGs, respectively (Table S2). Overall, a much higher number of DEGs (1496 downregulated and 1687 upregulated) were identified in the BC *Kdm6a*^*ΔP/-Y*^ clone, probably due to loss of all Y-linked genes in this clone, but this would need to be confirmed by analysis of additional clones with the same genotype (Figures 1B and S2E; Table S2). Nonetheless, there was concordance between the *Kdm6a*^*ΔP/-Y*^ clone and the *Kdm6a*^*Δ/Y*^ clones for 167/195 (86%) and 291/334 (87%) downregulated and upregulated DEGs, respectively (Figure S2E). In the single female BC clone (*Kdm6a*^*ΔP/ΔP*^) we found 263 downregulated and 203 upregulated DEGs (p-value < 0.1 and a fold-change cutoff of 1.5) (Figure 1C; Table S2). Consistent with our previous study, *Rhox6/9* were downregulated (1.4 fold or greater) in the female clone (Figure S2C) (Berletch et al., 2013). However, there was little overlap in DEGs between the BC female and male KO clones, possibly due to sex differences, clonal variation and/or batch effects (Figure S2E). Interestingly, more DEGs were in common between BC clones *Kdm6a*^*ΔP/-Y*^ and *Kdm6a*^*ΔP/ΔP*^, possibly reflecting the absence of Y-linked genes in both cell lines (Figure S2E).

**Figure 1:**
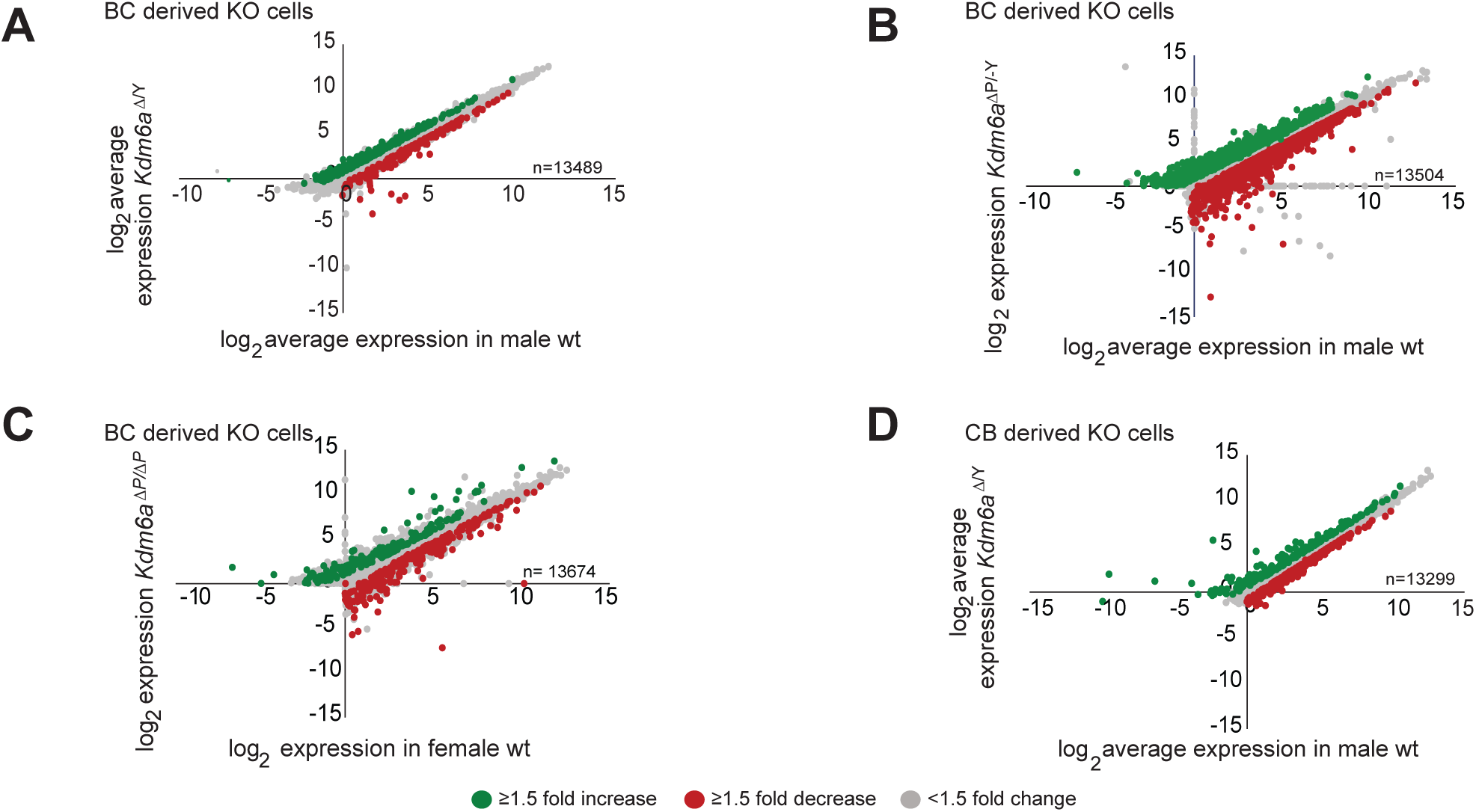
Global gene expression changes after KDM6A KO *(See also Figures S1-3)*. (**A-D**) Scatter plots of differential gene expression in three BC male *Kdm6a*^*Δ/Y*^ clones versus two male BC wt clones (**A**), one BC male *Kdm6a*^*ΔP/-Y*^ clone that has lost the Y chromosome versus two male BC wt clones (**B**), one BC female *Kdm6a*^*ΔP/ΔP*^ clone versus one female BC wt clone (**C**), and three CB male *Kdm6a* ^*Δ/Y*^ clones versus two CB male wt clones (**D**). Log_2_ values of genes with >1TPM in at least one replicate are shown. Genes labeled red are downregulated DEGs, in green, upregulated DEGs, and in grey are not considered differentially expressed. Only DEGs with a fold change ≥1.5 and FDR <0.05 or a p-value of <0.1 are shown.

In the CB cross we identified 334 downregulated and 213 upregulated DEGs in the KO *Kdm6a*^*Δ/Y*^ clones compared to wt clones (Figure 1D; Table S2). The lists of DEGs differ between the BC and CB crosses, with only 4 downregulated and 13 upregulated DEGs in common (Figure S2F; Table S2). When using a less stringent approach (≥1.25 log_2_ TPM fold change cutoff) there was a higher concordance between crosses, with 65/192 downregulated and 33/293 upregulated DEGs called in the BC cross showing a trend of decreased and increased expression in the CB cross, respectively, and another 35 downregulated genes in the BC KO clones showing lower TPM values in at least 2/3 CB KO clones (Table S2). When simply comparing the lists of genes with a ≥1.25-fold TPM value between male KO and wt clones in both crosses, 345 genes and 227 genes with decreased and increased expression, respectively, were shared between the reciprocal crosses (Figure S2F; Table S2). These genes include known KDM6A targets, e.g. *Bcar3, Foxh1*, and *Hsd17b11*.

Gene ontology (GO) analyses of downregulated DEGs in BC KO clones revealed developmental categories including embryo development (e.g. *Eomes, Dnmt3a*), mesoderm development (e.g. *T/Brachyury*), epithelial tube morphogenesis (e.g. *Lef1, Foxh1*), and tube closure (*Fzd8, Sall1)* (Figure 2A-C; Table S2) (Mi et al., 2019). GO terms for downregulated DEGs in CB KO clones include neurogenesis (e.g. *Sox6, Notch3*) and cell differentiation (e.g. *Pax6, Foxo6*), respectively (Figure 2D; Table S2). GO terms for genes that display a downregulation trend in common between the BC and CB crosses are enriched in developmental categories including neural tube development (e.g. *Foxh1*) and heart morphogenesis (e.g. *Pitx2*), consistent with neural tube defects and heart malformations observed *in utero* in *Kdm6a* KO mice (Figure 2E) (Shpargel et al., 2012; Wang et al., 2012). Interestingly, roof of mouth development and forebrain cell migration are also included (e.g. *Gdf11* and *Sox11*), consistent with high arched palate and other distinctive facial features observed in Kabuki syndrome caused by KDM6A mutations (Figure 2E, F) (Adam et al., 2019; Guo et al., 2018; Yap et al., 2020). Notably, there was decreased expression of *Nes*, a KDM6A target gene identified in neural crest cells, a type of cells that migrate ventrally and contribute to patterning and formation of all anterior facial bone and cartilage (Table S2) (Shpargel et al., 2017). Upregulated DEGs in BC and CB KO clones include developmental processes (e.g. *Prdm14, Klf4*) and brain development and a subset of genes known to be repressed by KDM6A (e.g. *Fam114a1, H1f0, Naglu*) (e.g. *Sox21, Wnt7b, Synj1*) (Figure 2A-E; Table S2) (Gozdecka et al., 2018).

**Figure 2:**
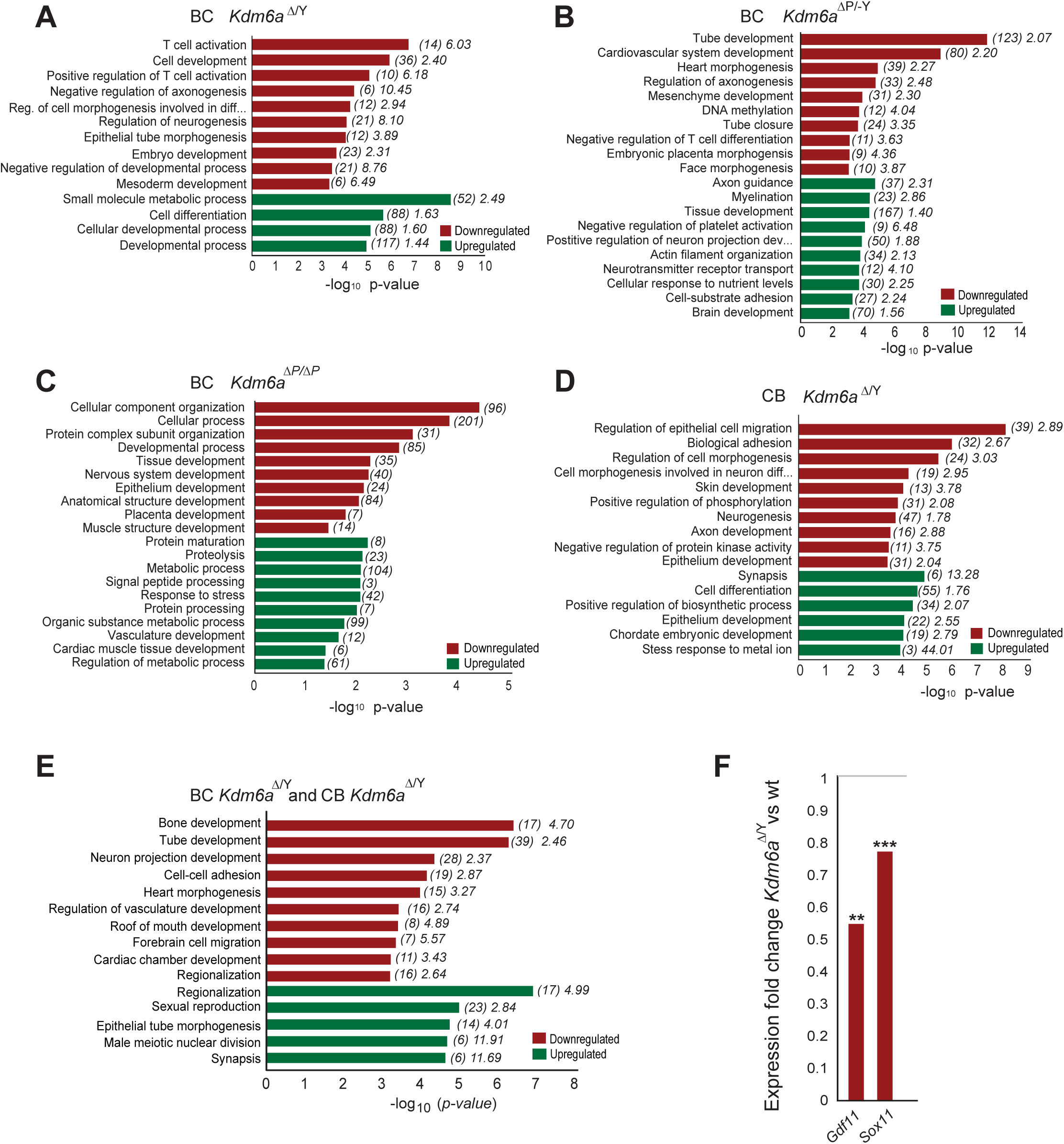
Gene ontology analysis. **A-D**. Gene ontology analysis for dysregulated DEGs in three BC male *Kdm6a*^*Δ/Y*^ clones (**A**), one BC male *Kdm6a*^*ΔP/-Y*^ clone that has lost the Y chromosome, (**B**), one BC female *Kdm6a*^*ΔP/ΔP*^ clone (**C**) and three CB male *Kdm6a* ^*Δ/Y*^ clones (**D**). Numbers in parentheses denote the number of genes included in the associated biological process, while those not in parentheses represent the fold enrichment over expected number of genes within a given biological process. All Y-linked genes were excluded from expression analysis in *Kdm6a*^*ΔP/-Y*^. (**E**) GO analysis of dysregulated genes common between male KO cells derived from BC and CB crosses. Numbers in parentheses represent the number genes in each category, while those not in parentheses represent the fold enrichment over expected number of genes within a given biological process. All processes are ranked by frequency in -log_10_ p-value. (**F**) Expression changes of *Gdf11* and *Sox11*, two genes known to be associated with orofacial and palate development, show downregulation both in KO cells from BC and CB crosses (**p ≤ 0.01). Fold decrease in TPM is shown.

Importantly, KDM6A KO did not lead to reduced expression of pluripotency and self-renewal genes, confirming that KDM6A is not necessary to maintain pluripotency nor did the editing process induce ES cell differentiation (Figure S2G) (Mansour et al., 2012). However, consistent with previous reports, differentiation kinetics for clone *Kdm6a*^*ΔE1*^ compared to wt ES cells showed slower formation of embryoid bodies and delayed appearance of neuronal cells after retinoic acid treatment (Figure S2H) (Morales Torres et al., 2013; Wang et al., 2012). Concordantly, KDM6A KO in all BC lines resulted in dysregulation of genes heavily involved in developmental processes.

Taken together, our findings support a role for KDM6A in the regulation of a subset of key developmentally critical genes in both male and female mouse ES cells. Despite cell line and clonal variability, downregulated DEGs affirm the role of KDM6A as an enhancer of gene expression, probably via its demethylase-dependent and -independent activities, whereas upregulated DEGs indicate a repressive function. We cannot rule out indirect effects on gene expression. Interestingly, some of the dysregulated genes common to reciprocal crosses could contribute to clinical features present in Kabuki type 2 syndrome.

### The majority of gene expression changes in KDM6A KO ES cells is allele-specific

To measure allelic gene expression, we assigned RNA-seq reads to parental alleles using SNPs between C57BL6/J and *M. castaneus* in the hybrid F1 mouse ES cells from the BC and CB crosses. Due to a lack of informative SNPs, about 6% of genes could be not be classified as having either bi-allelic expression (diploid) or parent-of-origin allelic expression bias. Allele-specific differential gene expression changes were identified using DESeq2 with similar parameters as described above (log_2_ fold change ≥1.5 fold, FDR< 0.05). Scatter plots of differential expression between KDM6A KO and wt clones considering maternal and paternal alleles are similar to those generated using diploid data (Figures S3A-D). However, allelic analyses were done independently from the diploid analyses, resulting in a few differences between the allelic and diploid DEG list (see methods).

Surprisingly, allelic expression changes following KDM6A KO in male BC clones *Kdm6a*^*Δ/Y*^ and *Kdm6a*^*ΔP/Y*^ versus wt showed that the majority of downregulated (167/194) and upregulated (139/189) DEGs with informative SNPs demonstrate changes limited to one allele (Table 1). This was also observed in male CB clones *Kdm6a*^*Δ/Y*^, with 237/269 and 171/198 genes demonstrating down-and upregulation limited to one allele, respectively (Table 1). Downregulated DEGs in KO clones were categorized into three groups: group A includes genes downregulated on the maternal allele, group B, on the paternal allele, and group C, on both alleles (Figure 3A, B; Table 1). Upregulated DEGs in KDM6A KO clones were categorized into three groups: group D includes genes upregulated on the maternal allele, group E, on the paternal allele, and group F, genes on both alleles (Figure 3C, D; Table 1).

**Table 1:** The number of genes in each group (see text) following KDM6A KO is shown. BC and CB DEGs refer to differentially expressed genes between wt and CRISPR KO clones from reciprocal crosses. See also Figure 3.

| *Allele* | *Group* | *Expression change* | *BC DEGs* | *CB DEGs* |
| --- | --- | --- | --- | --- |
| Maternal | A | Downregulated | 109 | 133 |
| Paternal | B | Downregulated | 58 | 104 |
| Biallelic | C | Downregulated | 27 | 32 |
| Maternal | D | Upregulated | 68 | 109 |
| Paternal | E | Upregulated | 71 | 62 |
| Biallelic | F | Upregulated | 50 | 27 |

**Figure 3:**
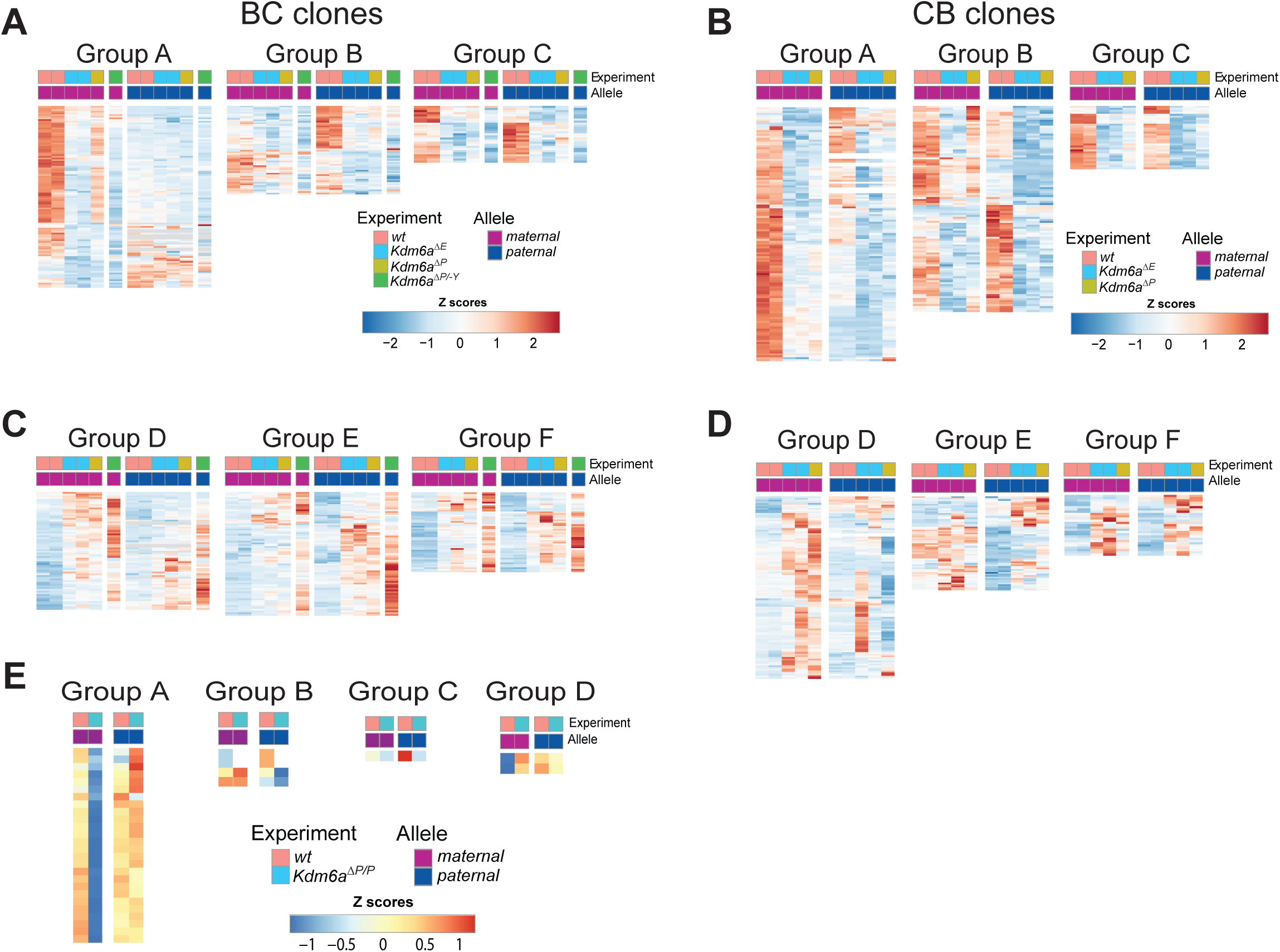
Allele-specific gene expression changes after KDM6A KO *(See also Figure S4)*. (**A**) Heatmaps of normalized allelic gene expression show the extent of fold changes for maternal alleles (left 6 columns of boxes) and paternal alleles (right 6 columns of boxes) of downregulated DEGs (gene names listed at right) in two male wt controls, three male BC KO (*Kdm6a*^*Δ/Y*^), including clones *Kdm6a*^*ΔE1*^ *Kdm6a*^*ΔE3*^, and *Kdm6a*^*ΔP4*^. Data for BC clone *Kdm6a*^*ΔP/-Y*^ (*Kdm6a*^*ΔP6*^) are shown in the 6th separate column. Z values (color-coded) representing deviations from the mean for each row are shown for groups A, B, C (see text). X-linked gene expression from the paternal allele was masked as zero (white boxes). See also Table S3. (**B**) Same analysis as in A for CB clones. (**C**) Same analysis as in A for upregulated DEGs in groups D, E and F (see text). (**D**). Same analyses as in C for CB clones. (**E**) Allelic analysis of downregulated (groups A, B, C) and upregulated DEGs (only group D genes were identified) in the female BC *Kdm6a*^*ΔP/ΔP*^ clone versus wt. Same analysis as in A and C.

Considering downregulated genes in the BC cross group A comprises 109 genes, whereas group B consists of only of 58 genes, suggesting preferential downregulation of maternal alleles (Figure 3A; Table 1). Many DEGs in group A are involved in immune functions (e.g. *Id1, Trim12c*), neurological development (e.g. *Arhgef9, Ddhd2*), placental and fetal development (e.g. *Phlda2, Meg3*), or are highly expressed in placenta (e.g. *Nip7, Lsm4, Xlr3c, Xlr5a, Xlr5c*), whereas group B genes include neurodevelopmental regulators (e.g. *Sox11, Foxd3*) (Table S3). The three male BC *Kdm6a*^*Δ/Y*^ clones and the one BC *Kdm6a*^*ΔP/-Y*^ clone had a similar pattern of downregulation from maternal alleles, indicating no Y chromosome effect on allelic expression changes after KDM6A KO (Figure 3A; Table S3). Despite a lack of statistical power due to the availability of a single female BC clone *Kdm6a*^*ΔP/ΔP*^ more genes were downregulated from the maternal than the paternal allele in this clone (26 and 4, respectively) (Figure 3E). Among downregulated genes in BC clones we found a subset of maternally but not paternally expressed imprinted genes. These maternally expressed imprinted genes include those in the *Meg3* cluster, *Xlr* cluster genes, and *Phlda2* (Figures S4A-D; Table S3). In addition, *H19* showed a significant decrease in clones *Kdm6a*^*ΔE1*^ and *Kdm6a*^*ΔE3*^ (*p=*.*01)* and a trend for a decrease in clones *Kdm6a*^*ΔP4*^ and *Kdm6a*^*ΔP6*^ (Figure S5D). A corresponding increase was observed for *Igf2* in *Kdm6a*^*ΔE1*^ (Figure S5F). Importantly, the maternal allele bias in downregulated genes in BC KO clones was maintained in CB KO clones, with 133 group A genes and 104 group B genes, respectively (Figure 3B; Table 1). However, there was little overlap between allelic DEGs categorized into the groups described above (Table S3). Nonetheless, some biological processes enriched for maternally downregulated genes in BC and CB KO clones were in common, e.g. placenta development (e.g. *Cdx4, Met*). Considering trends in expression based on TPM values, 38/100 (38%) group A and 27/46 (59%) group B genes identified in male BC clones showed a trend of decreased expression from the maternal and paternal allele in male CB clones, respectively (Table S4). Changes in maternally expressed imprinted genes observed in the BC clones were not verified in the reciprocal CB cross where there were few changes in imprinted gene expression, suggesting a clonal or strain-specific effect (Figure S4E).

In contrast to downregulated genes, upregulated genes in BC clones did not display an allelic bias, as reflected by a similar number of genes in groups D (68) and E (71) in the male BC clones, but did display an allelic bias in groups D (109) and E (69) in the male CB clones (Figure 3C, D; Table 1). Included in group D from BC clones are several genes associated with reproduction (e.g. *Id4, Tead4*) and development (e.g. *Stx2, Rb1*), whereas group E includes genes involved in metabolic processes (e.g. *Pdk2, Apoe, Dcxr*). The majority of upregulated genes (60%) in the *Kdm6a*^*ΔP/-Y*^ clone showed a stronger increase in expression for all three groups D, E, F compared to *Kdm6a*^*Δ/Y*^ clones, suggesting widespread effects due to loss of Y-linked genes (Figure 3C). In the female *Kdm6a*^*ΔP/ΔP*^ clone only two maternally upregulated DEGs and no paternally upregulated DEGs were identified, again due to low statistical power (Figure 3E). Further comparisons between BC and CB clones were made to address allelic bias of gene expression in wt cells. A majority of group A genes identified in male BC clones (76/100 or 76%, excluding X-linked genes) and male CB clones (65/100 or 60%, excluding X-linked genes) had maternally biased expression in wt clones (Figure 3A, B; Table S4). This maternal allele biased expression of group A genes in wt was confirmed in an independently derived CB ES cell line (Marks et al., 2015). Group B genes showed a lesser paternal allele bias in wt clones, and upregulated groups did not exhibit an allele bias in expression in wt cells.

We conclude that KDM6A facilitates gene expression from one allele or the other with a preference for maternal alleles of a subset of mouse genes. Importantly, this preference is seen in KDM6A KO cells from both BC and CB crosses, suggesting a general role for KDM6A in control of maternal alleles. However, different genes are affected, suggesting that this role is not gene-specific. There may also be strain and clonal effects. We also find evidence of gene repression by KDM6A, which is amplified in cells lacking the Y-linked paralog *Uty*, possibly reflecting a complete loss of a repressive function shared by the paralogs.

### KDM6A regulates maternal and paternal alleles by distinct mechanisms

KDM6A is a demethylase that removes methylation at H3K27me3, but it is also thought to have other functions. To determine whether KDM6A regulates allelic gene expression by demethylation-dependent or -independent mechanisms, we investigated allelic chromatin accessibility and histone H3K27me3 changes in male ES cells after KDM6A KO in the BC cross. Allelic ATAC-seq analysis was done to compare clone *Kdm6A*^*ΔE1*^ and wt male ES cells at or near promoters of DEGs (±2 kb from the TSS). For genes specifically downregulated (group A) and upregulated (group D) on maternal alleles the expected decrease and increase in chromatin accessibility was observed, respectively (Figure 4A). In contrast, genes downregulated at paternal alleles (group B) did not exhibit a loss of chromatin accessibility, while upregulated genes (group E) showed the expected increase in chromatin accessibility (Figure 4B). Comparisons of changes in ATAC-seq coverage between maternal and paternal alleles confirmed that among the three groups (A, B, C) of downregulated genes only group A showed the expected loss of accessibility, while all three groups (D, E, F) of upregulated genes show a gain in accessibility (Figures 4C-F; Table S5). Furthermore, levels of allelic chromatin accessibility were consistent with expression changes of KDM6A target genes (Figure 4). Notably, these differences in chromatin accessibility were observed only at promoters of DEGs, whereas there were no overall differences in accessibility levels at enhancers genome-wide regardless of parental allele (*p=0*.*06*, maternal allele; *p=0*.*56*, paternal allele) (Figure S5A). This is probably due to the relatively small number of KDM6A targets with expression changes following KO. However, chromatin accessibility is slightly increased at the promoters (Figure S5B), which does not appear to affect overall gene expression (Figure S3A).

**Figure 4:**
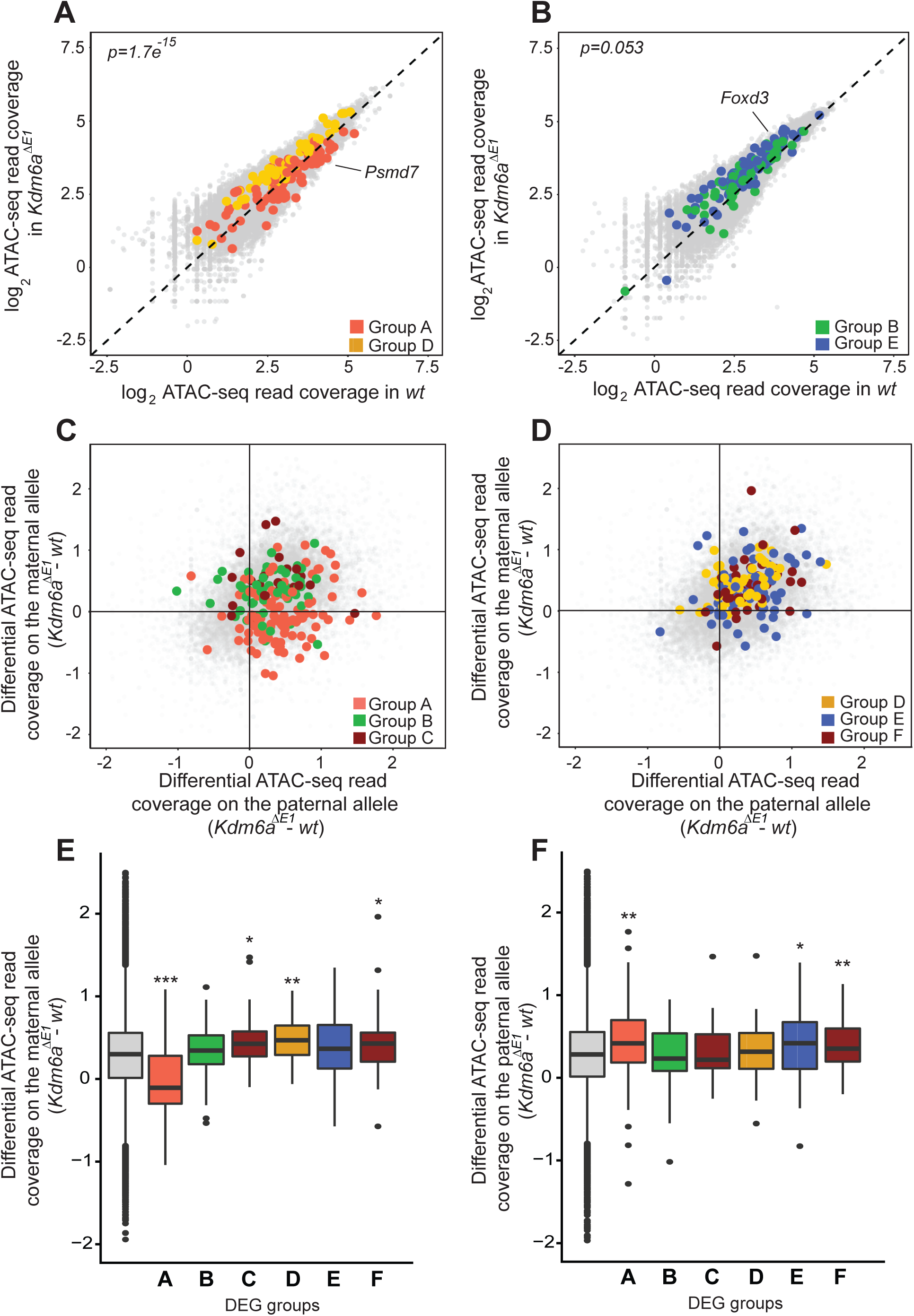
Reduced chromatin accessibility at maternally but not paternally downregulated genes after KDM6A KO *(See also Figures S5 and S6)*. (**A, B**) Scatter plots of ATAC-seq read coverage in log_2_ scale on maternal (**A**) and paternal (**B**) alleles in male BC *Kdm6a*^*ΔE1*^ cells versus wt. Allele-specific ATAC-seq reads around promoters (±2kb of the TSS) were used to calculate promoter coverage normalized by the background (genomic regions 5kb away from any ATAC-seq peaks) level. (**A**) Maternally downregulated genes (group A) show a shift below the diagonal (*p=1*.*7*^*e-15*^), indicating loss of DNA accessibility on the maternal allele in *Kdm6a*^*ΔE1*^ cells, while upregulated genes (group D) show a shift above the diagonal due to increased accessibility. (**B**) Paternally downregulated genes (group B) and upregulated genes (group E) both show a shift above the diagonal, indicating increased accessibility (*p=0*.*053*). (**C, D**) Scatter plots of ratios of ATAC-seq read coverage between clone *Kdm6a*^*ΔE1*^ and wt cells at promoters (±2kb of the TSS) on maternal alleles versus paternal alleles. Downregulated DEGs (**C**) (color-coded for genes in groups A, B, C) in group A are in the lower right quadrant, but, surprisingly, those in groups B and C are in the upper right quadrant. Upregulated DEGs (**D**) (color-coded for genes in groups D, E, F) are mostly in the upper right quadrant. (**E, F**) Box plots of average ratios of ATAC-seq read coverage between clone *Kdm6a*^*ΔE1*^ and wt cells at promoters (±2kb of the TSS) of maternal alleles (**E**) and paternal alleles (**F**) of downregulated DEGs (color coded for groups A, B, C; total genes in grey) and upregulated DEGs (color-coded for groups D, E, F, total genes in grey). Group A genes show a significant change in DNA accessibility, which is concordant with downregulation or upregulation (\*\*\**p=4*.*1*^*e-10*^ *and **p=9*.*7*^*e-07*^). No such correlation appears for group B genes. Group E genes show a minor increase in accessibility on the paternal allele (group E; \**p=0*.*012*), and group F genes on both alleles (group F; *\*p=0*.*02* and *\*\*p=0*.*006)*.

Next, we performed allele-specific ChIP-seq for H3K27me3, which demonstrated a global increase across the genome in clone *Kdm6a*^*ΔE1*^ compared to wt ES cells, with peak density higher on both alleles, consistent with the role of KDM6A as a demethylase (Figure 5A). Profiling of the control *Hoxb* cluster confirmed an increase in H3K27me3 enrichment in KDM6A KO cells and cells with KDM6A demethylase activity, as previously reported (Figure S5C) (Shpargel et al., 2017; Shpargel et al., 2014; Yu et al., 2019). A significant increase in H3K27me3 was observed at the promoters of paternally and bi-allelicly downregulated genes (groups B and C), but not at those of maternally downregulated genes (group A) (Figures 5B-D). Surprisingly, an increase in H3K27me3 was also observed at promoters of upregulated DEGs (groups D, E, and F), consistent with other reports that suggest that increased H3K27me3 levels do not always correlate with decreased gene expression (Figures 5B-D) (Gentile et al., 2019; Shpargel et al., 2017; Young et al., 2011). To confirm that the changes in H3K27me3 levels observed were due to loss of KDM6A rather than to changes in levels of PRC2 components, we verified the lack of significant changes in expression of genes encoding PRC2 complex members (*Ezh1, Ezh2, Suz12, Eed, Rppb7*) (Figure S5G). Note that this analysis does not rule out a disruption in the recruitment of these proteins.

**Figure 5.**
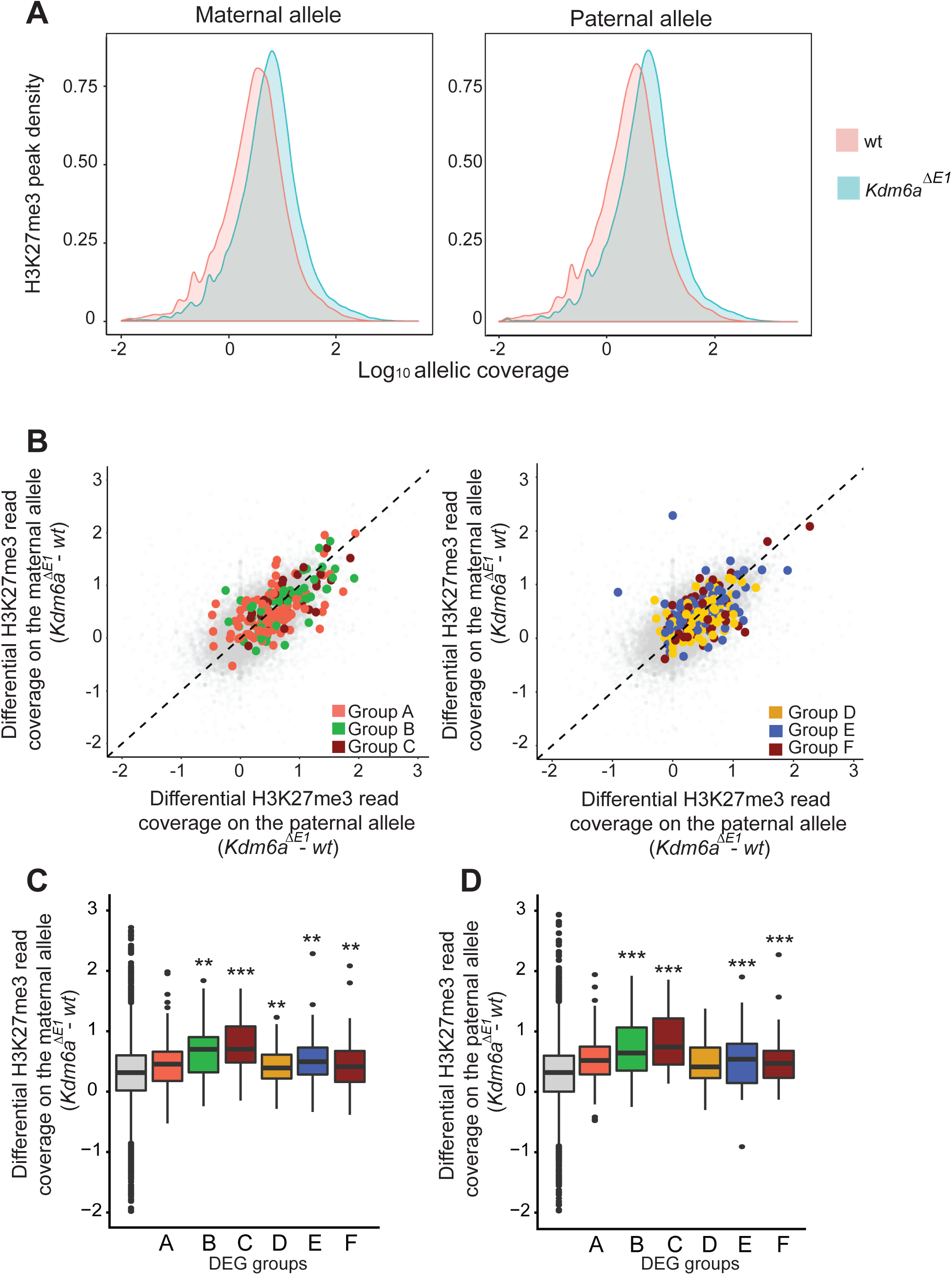
Allelic changes in H3K27me3 after KDM6A KO *(See also Figures S5 and S6)*. (**A**) Allelic H3K27me3 peak densities in wt and *Kdm6a*^*ΔE1*^ cells show an increase in H3K27me3 across the genome as measured by a shift in allelic peak density on the maternal and paternal alleles (*p=2*.*2e*^*-16*^). (**B**) Scatter plots of ratios of H3K27me3 ChIP-seq read coverage between clone *Kdm6a*^*ΔE1*^ and wt cells at promoters (±2kb of the TSS) on maternal alleles versus paternal alleles of downregulated DEGs (color-coded for genes in groups A, B, C) and upregulated DEGs (color-coded for genes in groups D, E, F). Values were normalized to the input and converted to log scale. (**C, D**) Box plots of average ratios of H3K27me3 ChIP-seq read coverage between clone *Kdm6a*^*ΔE1*^ and wt cells at promoters (±2kb of the TSS) on maternal alleles (**C**) and paternal alleles (**D**) of downregulated DEGs (color-coded for groups A, B and C, total genes in grey) and upregulated DEGs (color-coded for groups D, E and F, total genes in grey). Genes in groups B and C, but not in group A, show a significant correlated increase in H3K27me3 levels on both alleles (\*\**p=2*.*0*^*e-3*^ *and ***p=3*.*1*^*e-04*^). Genes in groups D, E, and F show a small but significant increase in H3K27me3 on maternal and paternal alleles (Groups D, E and F; \*\**p≤ 0*.*01 and ***p≤ 0*.*001*)

We then explored the relationship between allele-specific chromatin accessibility changes and H3K27me3 enrichment at promoters of DEGs in clone *Kdm6a*^*ΔE1*^ compared to wt ES cells. Decreased accessibility at maternally downregulated DEGs (group A) was rarely associated with increased levels of H3K27me3, e.g. at *Psmd7* (Figures 6A, C; Table S5). In contrast, paternally downregulated DEGs (group B), e.g. *Foxd3*, show a significant increase in H3K27me3 at their promoter, but no accompanying loss of chromatin accessibility (Figures 6B, C; Table S5). Bi-allelicly downregulated DEGs (group C), e.g. *Bcl11a*, behaved similarly to paternally downregulated DEGs (Figures 6A, C). Based on published H3K27ac profiles in wt ES cells, promoters of group A genes were significantly enriched in H3K27ac compared to group B genes, suggesting a potential demethylase-independent mechanism for KDM6A, which may involve acetylation of H3K27 on maternal alleles (Figure 6C; Figure S5F). We verified that expression levels of genes encoding chromatin remodeling proteins known to mediate H3K27 acetylation by association with KDM6A in the MLL complex (*Ep300, Smarca4, Eomes*) were unchanged in KO clones (Figure S5G). Again, this analysis does not exclude changes in recruitment of these proteins.

**Figure 6:**
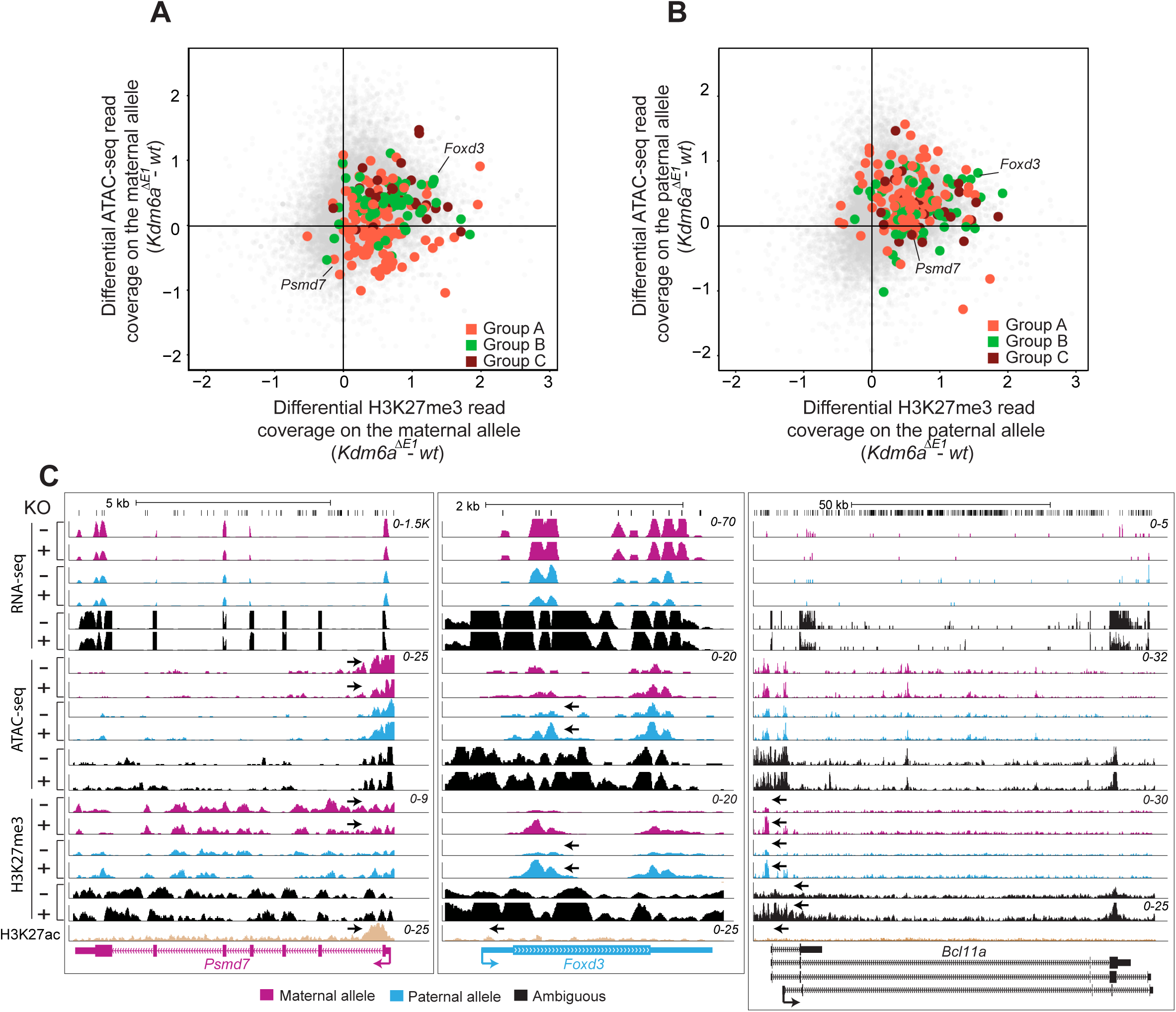
Epigenetic features of genes regulated by KDM6A differ depending on the parental allele *(See also Figure S5 and S6)*. (**A, B**) Scatter plots of ratios of ATAC-seq read coverage versus ratios of H3K27me3 ChIP-seq read coverage between clone *Kdm6a*^*ΔE1*^ and wt cells at promoter regions (±2kb of the TSS) of maternal (**A**) and paternal (**B**) alleles. A subset of maternally downregulated genes (group A) shows a significant decrease in DNA accessibility but no corresponding increase in H3K27me3 levels (\*\*\**p=4*.*1*^*e-10*^) on their maternal allele. In contrast, a subset of paternally downregulated genes (group B) shows a significant increase in H3K27me3 levels (*\*\*\*p=3*.*1*^*e-04*^) but no significant decrease in promoter DNA accessibility on the paternal allele. (**C**) UCSC genome browser (GRCm38/mm10) views of RNA-seq, ATAC-seq and H3K27me3 ChiP-seq profiles at the maternally downregulated gene *Psmd7*, the paternally downregulated gene *Foxd3*, and the bi-allelically downregulated gene *Bcl11a* in *Kdm6a*^*ΔE1*^ (KO+) and wt (KO-) cells. Also shown are published data for H3K27ac ChIP-seq in wt cells (Wang et al., 2017).

Our results point to KDM6A’s potential to regulate gene expression through histone demethylation-independent mechanisms on maternal alleles and histone demethylation-dependent mechanisms on paternal alleles of target genes. We propose a model in which group A genes are downregulated in KDM6A KO due to loss of accessibility, while group B and C genes are downregulated due to gain of H3K27me3 (Figure 7).

**Figure 7:**
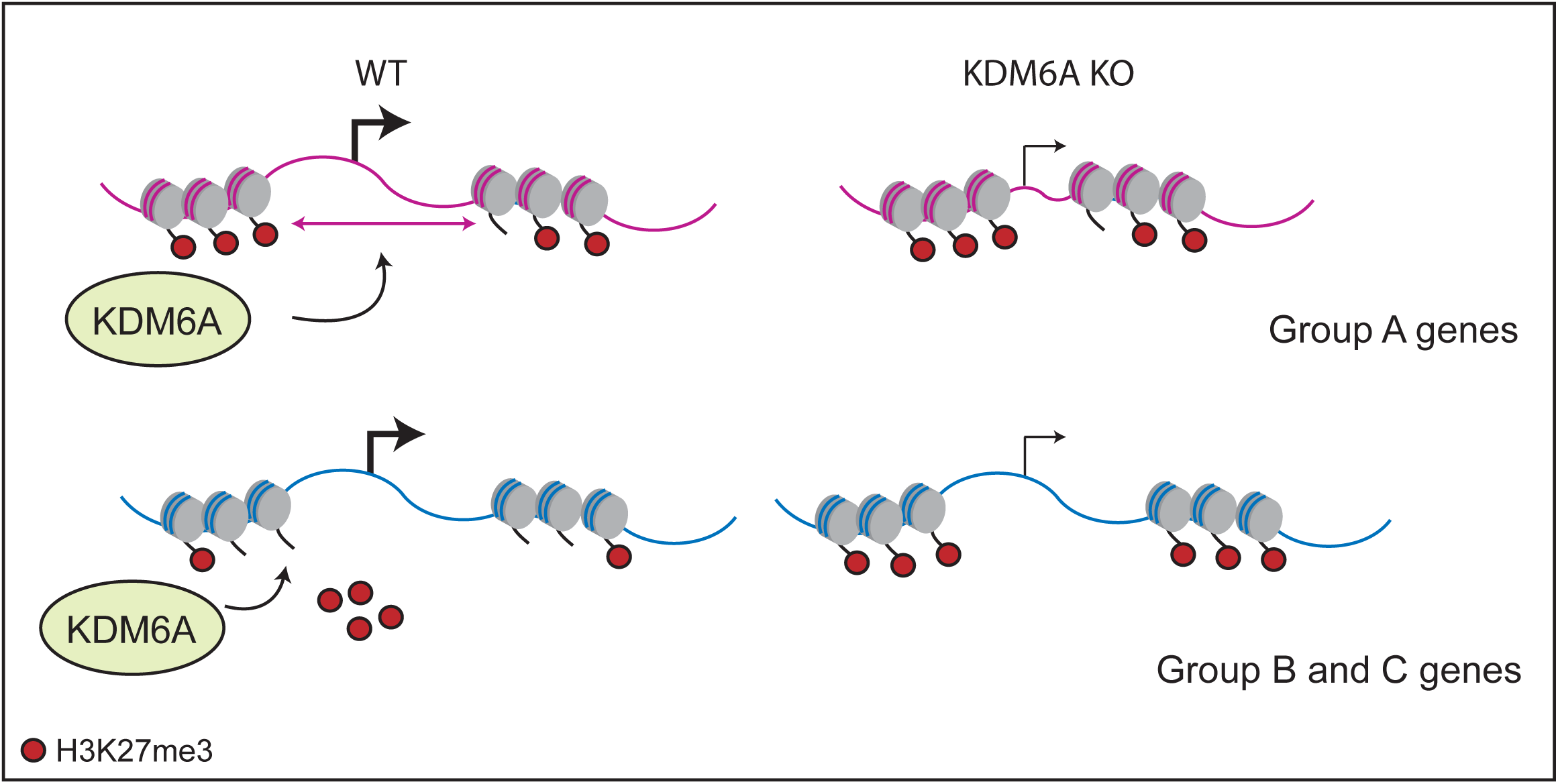
KDM6A regulates parental alleles of target genes through different mechanisms. Schematics of observed changes associated with downregulation of gene expression at maternal (pink) and paternal (blue) alleles depicts chromatin accessibility and H3K27me3 levels at groups A, B and C genes in wt and KDM6A KO clones. We propose that group A genes are regulated by KDM6A mainly via a demethylation-independent mechanism that enhances chromatin accessibility and gene expression without loss of H3K27me3, while groups B and C genes are mostly regulated by KDM6A via demethylation of H3K27me3 for increased gene expression.

## DISCUSSION

Here, we show that KDM6A KO in male and female ES cells derived from reciprocal crosses between C57BL/6J and *Mus castaneus* results in significant dysregulation of genes involved in development and reproduction. We also show that the majority of dysregulation occurs from one allele or the other. Interestingly, our data illustrate a maternal bias in downregulation in KO clones from both reciprocal crosses. Furthermore, a large subset of these targets also exhibited maternal expression bias in wt ES cells from reciprocal crosses. Finally, KDM6A appears to utilize distinct mechanisms to regulate maternal and paternal alleles.

Gene regulation by KDM6A mainly depends on its catalytic function. Indeed, depletion or inhibition of KDM6A results in decreased gene expression due to higher levels of the repressive histone mark H3K27me3 at promoters and/or enhancers of target genes (Berletch et al., 2013; Faralli et al., 2016; Mansour et al., 2012). Conversely, higher *KDM6A* expression in cancer cells causes a decrease of H3K27me3 enriched heterochromatin (Gurrion et al., 2017). Aside from its catalytic function KDM6A further enhances gene expression through histone demethylase-independent mechanisms that involve the recruitment of the MLL complex (Miller et al., 2010; Shpargel et al., 2017; Wang et al., 2017). Finally, KDM6A may also act as a repressor of gene expression, underscoring the complexity of this protein (Gozdecka et al., 2018; Lee et al., 2012; Miller et al., 2010; Shpargel et al., 2017).

We found that KDM6A KO in male and female mouse ES cells results in significant dysregulation of genes involved in development, immune and neuronal cell functions, metabolism, and reproduction. We and others have previously shown that genes necessary for germ layer development and ES cell differentiation are targets of KDM6A (Berletch et al., 2013; Morales Torres et al., 2013; Wang et al., 2012). It should be noted that indirect effects of downregulated or upregulated genes may contribute to widespread changes in expression. In fact, KDM6A’s primary role being to enhance gene expression via removal of a repressive histone mark, downregulated genes are more likely subject to direct effects, while upregulated genes following KDM6A KO are more likely subject to indirect effects. In particular, the more than 2-fold reduction in *Dnmt3a* expression observed in KDM6A KO male ES cells derived from the BC cross could cause DNA hypomethylation and explain the observed upregulation of DNMT3A target genes, such as *Celsr1* and *Fah* (Haney et al., 2016). Similarly, the decrease in *Tet1* expression in KO female ES cells could explain increased expression of TET1 target genes such as *Casp8*. Consistent with upregulation as an indirect effect of KDM6A KO a large portion of downregulated genes in KDM6A KO cells show a similar trend in BC and CB crosses, while the same is not true for upregulated genes.

KDM6A targets common to both reciprocal crosses include genes such as *Gdf11* and *Sox11* previously implicated in development of the roof of the mouth and hard palate, malformations of which are often seen in Kabuki syndrome due to mutations in *KDM6A* (Bogershausen et al., 2016; Silva-Andrade et al., 2019; Yap et al., 2020). *GDF11* mutations in human lead to orofacial abnormalities, while *SOX11* has a role in human palatogenesis (Cox et al., 2019; Khan et al., 2018; Suzuki et al., 2018). Our findings of additional dysregulated genes may help identify other gene targets implicated in congenital defects seen in Kabuki syndrome (Bogershausen et al., 2016; Lintas and Persico, 2018).

Unexpectedly, our allelic expression analyses showed that the majority of gene dysregulation occurred from one allele or the other with a maternal allele bias in KDM6A targets. Importantly, this maternal bias was maintained in both reciprocal crosses, even though different genes were dysregulated. Female-biased expression of *Kdm6a* due to its escape from X-inactivation is consistent with a role for KDM6A in differential regulation of maternal and paternal genomes during early development (Xu et al., 2008). Our previous studies also implicated KDM6A in the regulation of reproduction related *Hox* genes *Rhox6/9* (Berletch et al., 2013). Here we find that a number of genes with maternal allele-specific downregulation in KDM6A KO cells derived from the BC cross, including *Dmrt1, Cbfb*, and *Stox2*, are associated with reproductive processes (Fenstad et al., 2010; Matson et al., 2011; Tunster et al., 2016; Wilson et al., 2016). In addition, imprinted *Xlr* genes implicated in reproduction are downregulated in KDM6A KO cells, along with other maternally expressed imprinted genes, such as *Meg3*, a lncRNA gene that interacts with PRC2 components including JARID2 (da Rocha et al., 2014; Kaneko et al., 2014; Mondal et al., 2015; Yen et al., 2018). Among the few paternal allele-specific downregulated DEGs in cells derived from the BC cross, are *Fkbp6* and *Foxd3*, also implicated in reproduction. *Fkbp6* has a role in male fertility, with targeted inactivation resulting in azoospermic males (Crackower et al., 2003). *Foxd3* is necessary for proper maintenance of neural crest structures in mouse embryos, consistent with the observed lethality of female mouse embryos homozygous for a *Kdm6a* mutation (Teng et al., 2008). In KDM6A KO female ES cells derived from the BC cross several downregulated genes are also implicated in fertility and reproduction, but these genes differ from those found in KO male clones, suggesting sex differences in KDM6A targets. KDM6A KO mice have a sex-specific phenotype, with male mice runted but able to survive, while female mice die during embryogenesis (Shpargel et al., 2012). This has been attributed to a compensatory role by UTY in males. Since UTY has little or no demethylase activity this compensation probably functions via demethylase-independent mechanisms (Shpargel et al., 2012). Loss of this function could partly explain the marked increase in the number of DEGs in the KDM6A KO clone that lost the entire Y chromosome, but we cannot exclude effects of other Y-linked genes.

H3K27me3 is known to play a critical role in the repression of the maternal allele at non-canonical imprinted sites, loss of which, leads to loss of imprinting (Inoue et al., 2018). H3K27me3 levels are biased toward the maternal allele in mouse pre-implantation embryos and human morula and large H3K27me3 domains are established on the maternal allele during oogenesis which persist in the early pre-implantation embryo (Collombet et al., 2020; Zhang et al., 2019). Taken together, these data suggest proper levels of maternal H3K27me3 are critical for development. Thus, histone demethylation-independent mechanisms of gene regulation by KDM6A on the maternal allele may have arisen in order to protect silenced H3K27me3 domains from being re-activated through removal of H3K27me3. Indeed, our results strongly suggest that distinct epigenetic mechanisms operate on maternal and paternal alleles for regulation of gene expression by KDM6A (Figure 7). The relatively small number of significantly downregulated genes identified in KDM6A KO cells despite a genome-wide increase in H3K27me3 levels confirms a low concurrence of direct KDM6A targets with a histone demethylation phenotype (Shpargel et al., 2017; Young et al., 2011). Paradoxically, the presence of the PRC2 complex has been shown to facilitate contacts between promoters and distal elements at the *Hox* locus leading to a contradictory effect on gene expression (Gentile et al., 2019). The lack of concurrence we observed between loss of chromatin accessibility and H3K27me3 enrichment at maternal alleles of some downregulated genes could be caused in part by a low level of *Meg3*, which may hamper PRC2 recruitment (Mondal et al., 2015).

Downregulation of maternal alleles in KDM6A KO cells may reflect decreased levels of H3K27ac through histone demethylase-independent mechanisms (Carr et al., 2007; Wang et al., 2017). At genes downregulated from the paternal allele or genes with bi-allelic downregulation, expression appears to be primarily influenced by increased H3K27me3 levels without loss of chromatin accessibility, a pattern also reported in embryonic mouse brain (Wilken et al., 2015). Paternal allele-specific removal of H3K27me3 by KDM6A has also been illustrated in null peri-implantation embryos where reactivation of the paternal X chromosome is impaired (Borensztein et al., 2017). Further analysis of additional epigenetic changes following KDM6A KO, such as DNA methylation, may reveal more complex networks of gene regulation by KDM6A. Finally, the differential regulation of maternally and paternally expressed alleles by KDM6A may reflect fundamental differences in the three-dimensional structure of chromatin at the alleles as was reported for imprinted loci (Deng et al., 2015).

Here, we show for the first time a maternal bias in gene regulation by a gene that escapes X inactivation, a uniquely female phenomenon. Our results implicate KDM6A as a master regulator of fetal growth and development through the control of a subset of maternally biased and non-imprinted genes. While our studies are performed in ES cells where these developmental pathways may not be active, our data provide information on what pathways and lineages may be impacted following induction of differentiation. We propose that KDM6A can act through two distinct mechanisms to enhance expression of target genes depending on the parental allele. Further research is needed to determine whether association with different protein/RNA complexes confers KDM6A with particular functions at specific loci and how these complexes are capable of discriminating parental genomes. Inclusion of allelic analyses of gene regulation will enhance our understanding of parent-of-origin effects and sex-specific differences.

## ACKNOWLEDGEMENTS

We thank Di Kim Nguyen and Xinxian Deng (University of Washington) for consultation on this study and for critical reading of the manuscript, Giancarlo Bonora (University of Washington) for help with allele-specific mapping of ATAC-seq, and Charlie Lee (University of Washington) for his help with next-generation sequencing. Hybrid mouse ES cells used in this study were graciously donated by Joost Gribnau (Erasmus MC, University Medical Center). This work was supported by NIH grants GM046883 (CMD), GM113943 (CMD), MH105768 (JB) and NSF grant DBI-1751317 (WM).

## AUTHOR CONTRIBUTIONS

Conceptualization, J.B. and C.M.D.; Investigation, J.B., H.F., G.N.F, N.P.; Formal analysis, W.M.; Data Curation, W.F.; Writing – Original Draft, J.B. and C.M.D.; Writing – Review and Editing, J.B., W.M., G.N.F., and C.M.D.; Visualization, J.B., W.M., and C.M.D.; Supervision, J.B. and C.M.D.; Funding Acquisition, J.B. and C.M.D.

## DECLARATION OF INTERESTS

The authors declare no competing interests

## SUPPLEMENTARY MATERIAL

**Supplementary Table S1: Summary of clones**

**Supplementary Table S2: Diploid expression changes in reciprocal crosses**

**Supplementary Table S3: Allele-specific expression changes in reciprocal crosses**

**Supplementary Table S4: Allelic characteristics of BC genes in CB cells**

**Supplementary Table S5: Allelic epigenetic changes at promoters of Kdm6a target genes**

**Supplementary Table S6: SNP and RNA-seq mapping metrics in wt and KDM6A KO cells**

### Supplementary figure legends

**Figure S1: CRISPR/Cas9 editing of *Kdm6a* in BC and CB mouse ES cells Related to Figures 1-7**. (**A**). Schematic shows location of the exonic deletion (*Kdm6a*^*ΔE*^) that removed exons 2-4 of *Kdm6a* in male BC and CB ES cells. Exons are shown as vertical bars with location of the PCR and RT-PCR primers (color-coded arrows) used to confirm the deletion and measure expression. Above, images of gels after electrophoresis of PCR products (gDNA), RT-PCR products (cDNA), and Western blot (protein) confirm KDM6A KO in male BC clones (*Kdm6a*^*ΔE1*^, *Kdm6a*^*ΔE3*^) and CB clones (*Kdm6a*^*ΔE2*.*5*^, *Kdm6a*^*ΔE2*.*7*^) compared to wt (color-coding refers to primers shown on schematic). *Actb* was run as a control for PCR and RT-PCR and Ponceau S staining was used as loading control for the Western blot. Partial *Kdm6a* sequence obtained by Sanger sequencing is shown below the schematic as verification of deletions in all male *Kdm6a*^*ΔE*^ clones compared to wt. (**B**) Schematic shows location of the promoter region deletion (*Kdm6a*^*ΔP*^) made in male BC and CB ES cells. Exons are shown as vertical bars with location of the PCR and RT-PCR primers (color-coded arrows) used to confirm the deletion and measure expression. Above, images of gels after electrophoresis of PCR products (gDNA) and RT-PCR products (cDNA) confirm KDM6A KO in male BC clones (*Kdm6a*^*ΔP4*^, *Kdm6a*^*ΔP6*^) and CB clones (*Kdm6a*^*ΔP2*.*1*^) compared to wt. *Actb* was run as a control for PCR and RT-PCR. Partial *Kdm6a* sequence obtained by Sanger sequencing is shown below the schematic as verification of deletions in all *Kdm6a*^*ΔP*^ male ES cell clones and in a female BC ES cell (E8) clone *Kdm6A*^*ΔP/ΔP*^ with a homozygous deletion of *Kdm6a* promoter. (**C-D**) UCSC genome browser (GRCm38/mm10) views of RNA-seq profiles for (**C**) BC male and female wt, BC male clones *Kdm6a*^*ΔE*^ and *Kdm6a*^*ΔP*^, and BC female clone *Kdm6a*^*ΔP/ΔP*^, and (**D**) CB male wt, and CB male *Kdm6a*^*ΔE*^ and *Kdm6a*^*ΔP*^ clones. Note that a low level of 3’ end reads are present in clones with deletion of exons 2-4, although there is no evidence of protein (see A above). In male and female clones with deletion of the promoter there are no reads over *Kdm6a*, including no reads overlapping the small alternative *Kdm6a* transcript, confirming absence of expression and suggesting the absence of a cryptic promoter for the smaller annotated transcript. (**E**) Gel electrophoresis of PCR products of gDNA to test for the presence/absence of *Uty. Uty* is absent due to loss of the Y chromosome in BC clone *Kdm6a*^*ΔP6*^. *Actb* was used as a PCR control. All primer sequences and sgRNAs are available upon request. (**F**) Evidence of off-target effect in male BC clones *Kdm6a*^*ΔE1*^ and *Kdm6a*^*ΔE3*^. RNA-seq genome browser profiles in BC derived *wt* and KDM6A KO male clones show absence of *Pdha1* reads in *Kdm6a*^*ΔE*^ but not *Kdm6a*^*ΔP*^ clones. The inset shows PCR analysis for regions A, B and C amplified by primers indicated by colored arrows, which confirms an off-target deletion of part of *Sh3kbp1* and all of *Map3k15* and *Pdha1* in *Kdm6a*^*ΔE*^ clones. No other KO clones show this off-target deletion (data not shown). *Actb* was run as a control.

**Figure S2: Expression changes in BC and CB KDM6A KO ES cells. Related to Figures 1 and 2**. (**A**) Principal component analysis (PCA) based on diploid expression values for all transcribed genes from RNA-seq in wt cells and *Kdm6a* KO clones derived from BC/CB reciprocal crosses. For the BC cross two male wt, one female wt, three male KO (*Kdm6a*^*Δ/Y*^), one male KO with no Y chromosome *(Kdm6a*^*ΔP/-Y*^) and one female homozygous KO (*Kdm6a*^*ΔP/ΔP*^) are included, while for the CB cross two male wt and three male KO (*Kdm6a*^*Δ/Y*^) are included. (**B**) Hierarchal clustering of the lines described in A. The color scale represents the sample to sample distance. (**C**) Expression fold changes between male BC *Kdm6a*^*Δ/Y*^ clones and wt measured by RNA-seq confirm decreased expression of known KDM6A targets *Wt1, Bcar3, Foxh1, Dnmt3a, Sox3, Hsd17b11* and loss of *Kdm6a* expression. Expression fold changes between female BC *Kdm6a*^*ΔP/ΔP*^ and wt show decreased expression of *Rhox6* and *Rhox9*. Expression is based on diploid analysis. **FDR <0.001. (**D**) Expression fold changes between male BC *Kdm6a*^*Δ/Y*^ clones and wt measured by quantitative RT-PCR analysis confirm downregulation of *Vrtn, Bcar3*, and *Sall2* (*p<0.01 using a student’s t-test) (Table S2). Expression is normalized to *Actb*. (**E**) Venn diagrams of downregulated and upregulated DEGs in male BC *Kdm6a*^*Δ/Y*^, male BC *Kdm6a*^*ΔP/-Y*^, and female BC *Kdm6a*^*ΔP/ΔP*^ clones. (**F**) Venn diagrams of downregulated and upregulated DEGs in male BC and CB *Kdm6a*^*Δ/Y*^ and Venn diagrams for the same clones but listing number of genes downregulated or upregulated as measured by fold TPM differences. (**G**) Expression fold changes between BC derived *Kdm6a*^*Δ/Y*^ clones and wt measured by RNA-seq show no significant expression decrease in several pluripotent genes (*Nanog, Pou5f1, Sox2, Cd9, Stat3, Fut4*) (*\*\*p <0*.*01* using a student’s t-test). (**H**) Deficiencies in ES cell differentiation potential following KDM6A KO were tested by removal of LIF and by *all*-*trans* retinoic acid (RA) treatment. Smaller and less dense embryoid bodies were observed 6 days after removal of LIF in *Kdm6a*^*ΔE1*^. In the presence of RA, wt cells show morphological signs of differentiation after 8 days while *Kdm6a*^*ΔE1*^ cells remain similar in morphology to day 0 controls.

**Figure S3: Linear relationships between gene expression (diploid, maternal and paternal alleles) in wt and BC and CB KO clones. Related to figures 1 and 3**. (**A**-**D**) Scatter plots of average gene expression based on TPM values show a high correlation between wt and (**A**) the male BC *Kdm6a*^*Δ/Y*^ clones, (**B**) the male BC *Kdm6a*^*ΔP/-Y*^ clone, **(C)** the male CB *Kdm6a*^*Δ/Y*^, and (**D**) the female BC *Kdm6a*^*ΔP/ΔP*^ clone for diploid and allele-specific expression. Correlation coefficients are all ≥ 0.98.

**Figure S4. Expression of a subset of maternally expressed imprinted genes decreases after KDM6A KO in BC but not in CB clones. Related to figure 3**. (**A**) Scatter plot of average expression changes for canonical imprinted genes, either maternally (purple) or paternally (blue) expressed, in *Kdm6a*^*Δ/Y*^ (average from 3 clones) versus wt (average from 2 clones) based on diploid RNA-seq analysis in male BC derived clones. Only imprinted genes with >1TPM in at least one sample are shown. (**B**) RT-PCR analysis confirms decreased expression of *Meg3* and *Rian* in all four male BC KDM6A KO clones (*Kdm6a*^*ΔE1*^, *Kdm6a*^*ΔE3*^, *Kdm6a*^*ΔP4*^ and *Kdm6a*^*ΔP6*^), regardless to whether the Y chromosome is retained. CRISPR +/– labels indicate presence/absence of transfection with CRISPR/Cas9 and *Kdm6a* sgRNAs, with a + for *Kdm6a*^*ΔP*^ and *Kdm6a*^*ΔE*^ clones in which editing was successful. Non-edited CRISPR control clones (*Kdm6a*^*E14*^, *Kdm6a*^*P2*^) maintain expression of these genes. (**C**) UCSC Genome browser (GRCm38/mm10) view of allele-specific RNA-seq profiles at the maternally expressed *Dlk1*/*Mirg* polycistron region. Allelic read profiles on the maternal chromosome (purple), the paternal chromosome (blue) and diploid profiles (black) are based on SNP analysis in male BC *wt* clone (KO-), compared to male BC *Kdm6A*^*ΔE1*^ clone (KO+). SNPs are indicated at the top. *Mir* clusters A and B contain 11 and 39 annotated microRNAs, respectively. (**D**) UCSC Genome browser (GRCm38/mm10) view of allele-specific RNA-seq profiles at the three maternally expressed imprinted genes (*Xlr3c, Xlr5a, Xlr5c*). Allelic read profiles on the maternal chromosome (purple), the paternal chromosome (blue) and diploid profiles (black) are based on SNP analysis in male and female BC *wt* clones (KO-), compared to male BC *Kdm6A*^*ΔE1*^ clones and female *Kdm6a*^*ΔP/ΔP*^ clone (KO+), respectively. SNPs are indicated at the top. (**E**) Same analysis as in (**A**) for male CB KDM6A KO clones.

**Figure S5: Chromatin changes genes regulated by KDM6A. Related to Figures 4-5. (A)** Box plots of ATAC-seq read coverage differences between male BC clone *Kdm6a*^*ΔE1*^ and wt cells at enhancers. Diploid ATAC-seq coverage and allelic coverage are shown. Enhancers (n=25,346) were defined as regions enriched in H3K4me3 and H3K27ac in male ES cells as described (Wang et al., 2017). No significant chromatin accessibility difference was seen (*p=0*.*06*, maternal allele; *p=0*.*56*, paternal allele), nor was there a significant difference between alleles *(p=0*.*09)*. **(B)** Same analysis as in A but for promoters (n=20,745). ATAC-seq coverage was calculated using ±2kb surrounding the transcription start sites determined using GENCODE. There were significant increases in chromatin accessibility at promoters on both alleles (p<2.2^e-16^) using a one sample t-test. (**C**) UCSC Genome browser (GRCm38/mm10) view of H3K27me3 profiles in wt and *Kdm6A*^*ΔE1*^ confirms a general increase across the entire *Hoxb* locus following KDM6A KO. (**D**) Expression fold changes between *Kdm6a*^*Δ/Y*^ clones and wt measured by RNA-seq for genes encoding known chromatin modifying proteins in the PRC2 complex (*Ezh1/2, Suz12, Eed, Rbbp7*) and the MLL complex (*Ep300, Smarca4, Eomes*). (**E**) H3K27ac coverage at promoters (±2kb of the TSS) of maternally (group A) and paternally (group B) downregulated DEGs shows a significant enrichment at maternally downregulated genes (\*\*\**p=1*.*07*^*e-5*^). H3K27ac peaks analyzed from previously published data (Wang et al., 2017). (**F**) UCSC Genome browser (GRCm38/mm10) view of allele-specific RNA-seq, ATAC-seq and H3K27me3 ChIP-seq profiles generated in male *wt* (KO-) and *Kdm6A*^*ΔE1*^ (KO+) at the imprinted genes *H19* (maternally expressed) and *Igf2* (paternally expressed). Also shown is the H3K27ac profile from published ChIP-seq data in wt cells (Wang et al., 2017). Locations of SNPs are indicated at the top. The DMR (differentially methylated region) is denoted by a green bar. (**E**) TPM expression fold change measured by RNA-seq for *H19* and *Igf2* in *Kdm6A*^*ΔE1*^ versus wt. *H19* downregulation is concordant to an expected increase in expression of the paternally expressed imprinted *Igf2*.

## STAR Methods

### KEY RESOURCES TABLE

| REAGENT or RESOURCE | SOURCE | IDENTIFIER |
| --- | --- | --- |
| Antibodies |  |  |
| Anti-KDM6A | Bethyl Laboratories. Inc. | Cat. A302-374; RRID: AB_1907257 |
| Anti-H3K27me3 | Millipore | Cat. 07-449; RRID: AB_310624 |
| Bacterial and Virus Strains |  |  |
| Stable competent E.coli | New England Biolabs | Cat. C3040H |
| Biological Samples |  |  |
| Chemicals, Peptides, and Recombinant Proteins |  |  |
| SYBR Green PCR Master Mix | Roche | 4385610 |
| FastDigest Bpil | Thermo Fisher | FD1014 |
| FastAP | Thermo Fisher | EF0651 |
| T4 PNK | NEB | M0201S |
| Quick Ligase | NEB | M2200S |
| PlasmidSafe exonuclease | Epicenter | E3101K |
| Plasmid transfection media | Santa Cruz Biotech | sc-108062 |
| Ultra Cruz transfection reagent | Santa Cruz Biotech | sc-395739 |
| Ponceau S | Sigma-Aldrich | P-3504 |
| Fetal Bovine Serum, Premium | Hyclone | SH30396.03 |
| Penicillin streptomycin | Sigma-Aldrich | P0781 |
| Leukemia inhibitory factor | Millipore | ESG1107 |
| Non-essential amino acids | Gibco | 11140-050 |
| DMEM, high glucose, no L-glutamine | Sigma-Aldrich | D5796 |
| All-trans retinoic acid | Sigma-Aldrich | R2500 |
| Puromycin dihydrochloride | Sigma-Aldrich | P8833 |
| Formaldehyde | Macron Fine Chemicals | UN1198 |
| Protein A agarose beads | GE Healthcare | 17-5138-01 |
| Glycine | Sigma-Aldrich | G-7126 |
| cOmplete, Mini, EDTA-free Protease Inhibitor Cocktail | Millipore-Sigma | 11873580001 |
| Critical Commercial Assays |  |  |
| GoScript RT-PCR kit | Promega | A5000 |
| TruSeq RNA sample prep kit, V2 | Illumina, Inc | RS-122-2001 |
| TruSeq DNA sample prep kit | Illumina, Inc | FC-121-2001 |
| RNeasy Mini-prep kit | Qiagen, Inc | Cat. 74106 |
| Nextra DNA library prep kit | Illumina, Inc | FC-121-1030 |
| DNeasy Blood and Tissue kit | Qiagen, Inc | Cat. 69506 |
| BCA Protein Assay kit | ThermoFisher | Cat. 23227 |
| NE-PER protein isolation kit | ThermoFisher | Cat. 78833 |
| ECL Western Blotting Substrate | Pierce | Cat. 32106 |
| Deposited Data |  |  |
| Raw and analyzed data | This paper |  |
| Deposited data from (Wang et al., 2017) were downloaded from GEO | This paper | GEO: GSE97703 |
| Experimental Models: Cell Lines |  |  |
| Cast/BL6 hybrid stem cells | Gift from Dr. Joost Gribnau, Erasmus MC |  |
| BL6/Cast hybrid stem cells | Gift from Dr. Joost Gribnau, Erasmus MC | (Barakat et al., 2011) |
| Experimental Models: Organisms/Strains |  |  |
| Oligonucleotides |  |  |
| Primers for PCR, qPCR and sgRNA oligos available upon request. | This paper | N/A |
| Recombinant DNA |  |  |
| pPGKpuro | Addgene | Cat. 11349 |
| pX330-U6_Chimeric_BB_cBh-hspCas9 | Addgene | Cat. 42230 |
| UTX CRISPR/Cas9 KO plasmids | Santa Cruz Biotechnology | Cat. sc-423635 |
| pX330-KDM6A_promoter KO plasmids | This paper | N/A |
| Software and Algorithms |  |  |
| R | R Core Team, 2018 | https://www.r-project.org/ |
| DAVID 6.8 | (Huang da et al., 2009) | https://david.ncifcrf.gov |
| Panther Gene Ontology | (Mi et al., 2019) | http://www.pantherdb.org/ |
| ChopChop | (Labun et al., 2016) | https://chopchop.cbu.uib.no/ |
| Deseq2 | (Love et al., 2014) |  |
| Bowtie 2 | (Langmead and Salzberg, 2012) | http://bowtie-bio.sourceforge.net/bowtie2/index.shtml |
| Tophat 2 | (Kim et al., 2013) | https://ccb.jhu.edu/software/tophat/index.shtml |
| MACS 2 | (Zhang et al., 2008) | https://github.com/taoliu/MACS |
| Picard/v2.14.1 |  | http://broadinstitute.github.io/picard/ |
| Other |  |  |
| UVP Doc-ItLS | UVP, LLC | NA |
| Next-seq sequencer | Illumina | NA |
| ABI 7900HT | Applied Biosystems | NA |
| Misonix 3000 Sonicator | Misonix | NA |
| Motic AE2000 inverted microscope | Motic | NA |

#### Lead Contact and Materials Availability

Further information and requests for reagents and resources, including high throughput sequencing raw data, and KO cell lines should be directed to and will be fulfilled by the Lead Contact, Joel B. Berletch.

### EXPERIMENTAL MODEL AND SUBJECT DETAILS

#### Cell lines

BC male (E14) and female (E8) hybrid ES cells as well as CB hybrid ES cells derived from reciprocal crosses between C57BL/6J and *Mus castaneus* were provided by J. Gribnau, Erasmus MC Rotterdam, NL. All ES cells were maintained in the presence of 1000 U/ml leukemia inhibitory factor (LIF) (Millipore) on a monolayer of chemically inactivated mouse embryonic fibroblasts (MEFs) in ES media containing high glucose DMEM media supplemented with 15% fetal bovine serum (FBS), 1% non-essential amino acids, 10 mg/ml APS, 0.1mM 2-mercaptoethanol and 25mM L-glutamine. ES cells were grown in a humidified incubator at 37°C and 5% CO_2_. Just prior to use, plates were enriched for ES cells by incubation on 0.1% gelatin coated dishes for 1h to allow MEFs to attach, followed by transfer to fresh gelatin-coated plates for overnight culture. Following expansion, ES cells were split once (1:10) to further reduce potential MEF contamination.

### METHOD DETAILS

#### ES cell differentiation

ES cells were differentiated as follows. For embryoid body (EB) formation, 2×10^6^ wt control and *Kdm6a*^*ΔE1*^ cells were cultured on non-adherent bacterial culture dishes without leukemia inhibitory factor (LIF) for 6 days. Media was changed every two days. For retinoic acid induced neuronal differentiation, 1.2×10^6^ wt control and *Kdm6a*^*ΔE1*^ cells were grown on gelatin-coated tissue culture plates without LIF and in the presence of 1μM all-trans retinoic acid (Sigma) for 8 days. Media was changed every two days. Pictures were taken at the indicated time point using a Motic AE2000 camera.

#### CRISPR/Cas9 gene editing

We chose a dual sgRNA approach because of the capacity to create large deletions (Byrne et al., 2015). In addition, two independent sets of sgRNAs were used to account for off target effects. First, plasmids containing sgRNAs (Santa Cruz Biotechnology) located in exons 2 and 4 were used to delete ∼45kb, resulting in multiple stop codons in five of the six possible translation frames. Six-Frame translation analysis was used for *in silico* verification of newly acquired translation stop codons following *Kdm6a* targeting. For a second CRISPR/Cas9 editing approach, we targeted the *Kdm6a* promoter region including exons 1 and 2 using sgRNAs designed using the CHOPCHOP online tool (Labun et al., 2016; Montague et al., 2014) (Figure S1B). sgRNAs were cloned into the CRISPR/Cas9 plasmid pX330 as described (Cong et al., 2013).

ES cells were transfected with sgRNAs containing CRISPR/Cas9 constructs and a plasmid carrying puromycin resistance (Addgene #11349) using UltraCruz® transfection reagent at a 3:1 CRISPR to pPGKpuro ratio in media with no antibiotics. Two days later, cells were selected for 48-72h with 1μg/ml puromycin in normal ES media. Cells were allowed to recover in media with antibiotic. Following recovery, cells were cloned into 96 well plates using serial dilutions. Clones were expanded and screened for deletions using PCR and Sanger sequencing. PCR analysis confirmed correct gene editing in two BC male ES cell clones (*Kdm6a*^*ΔE1*^, *Kdm6a*^*ΔE3*^) and two CB male ES cell clones (*Kdm6a*^*ΔE2*.*5*^, *Kdm6a*^*ΔE2*.*7*^) (Figure S1A). Sanger sequencing showed that in contrast to the CB KO clones, deletion breakpoints in the *Kdm6a*^*ΔE1*^ and *Kdm6a*^*ΔE3*^ BC clones were identical, suggesting that they were derived from a single targeting event (Figure S1A). PCR amplification and Sanger sequencing confirmed the predicted ∼3.7 kb deletion in two BC male ES cell clones (*Kdm6a*^*ΔP4*^, *Kdm6a*^*ΔP6*^) and one CB ES clone (*Kdm6a*^*ΔP2*.*1*^) (Figure S1B). Additionally, we confirmed homozygous promoter editing in one BC female ES cell clone (*Kdm6a*^*ΔP/ΔP*^) (Figure S1B). *Kdm6a* expression in *Kdm6a*^*ΔE*^ and *Kdm6a*^*ΔP*^ clones was assayed by RT-PCR using primers specific for regions inside the deletion (spanning exons 3-6) and regions downstream of the deletion (spanning exons 23-26). RT-PCR and RNA-seq analyses showed low-level expression of exons downstream of the deletion, but Western blot analysis showed complete ablation of the KDM6A, consistent with premature stop codons and published *Kdm6a* exon KO models (Figure S1A) (Shpargel et al., 2012). RNA-seq and RT-PCR expression analyses verified the lack of *Kdm6a* expression in clones with a promoter deletion (Figures S1B-D). Note that the small alternative transcript annotated in UCSC (http://genome.ucsc.edu/) was absent in these clones as shown by the lack of reads in this region (Figures S1C and S1D). We determined that *Uty* was absent in one of the BC male clones with a *Kdm6a* promoter deletion (*Kdm6a*^*ΔP6*^) due to loss of the Y chromosome (Figure S1E). All sgRNAs are listed in Table S1.

#### Cell lysate preparation

Nuclear protein lysates were prepared using the NE-PER protein extraction kit according to manufacturer’s instructions. All plasmid preparations were made using either the Qiagen Mini-prep or Maxi-prep kits according to manufacturer’s instructions. All RNA and DNA lysates were prepared using Qiagen RNeasy mini kit and the Qiagen DNeasy blood and tissue kit, respectively.

#### Reverse-transcription quantitative PCR

RNA was extracted from cells as described above followed by cDNA synthesis with the GoScript reverse transcription kit. Relative transcript levels were determined using SYBR Green PCR master mix on the ABI 7900HT machine. All qRT-PCR assays were conducted in triplicates and normalized to *Actb*; the comparative CT method was used to analyze the data. All primer sequences are listed in Table S1.

#### Western blotting

Protein immunoblots were done to confirm protein ablation in *Kdm6a*^*ΔE*^ CRISPR deletions. Nuclear protein was extracted from *Kdm6a*^*ΔE1*^ and *Kdm6a*^*ΔE3*^ KO clones as well as control cells as described above. Concentrations were determined using the BCA protein assay kit. Equal concentrations of protein lysates were loaded onto SDS-PAGE gels, Tris-glycine 8%. Proteins were transferred to nitrocellulose membranes in Tris-glycine plus methanol buffer overnight at 4°C. After blocking After blocking with Tris-buffered saline (TBS), 0.1% Tween-20, 5% nonfat dry milk for one hour at room temperature, membranes were incubated at 4°C overnight with the primary antibody: rabbit polyclonal antibody against KDM6A (1:3000). After primary antibody incubation, immunoblots were incubated with HRP-conjugated secondary antibodies (1:10000) and visualized by chemiluminescence. Ponceau S staining of total protein served as a loading control.

#### Sanger sequencing

All plasmid, DNA, and RNA lysates used for Sanger sequencing were made using the methods described above. All reactions were prepared and submitted according to the protocols provided by Eurofin Genomics (www.operon.com).

#### RNA-seq with diploid analysis

RNA-seq indexed libraries were prepared using Illumina TruSeq RNA sample preparation kit V2. For BC derived ES cells libraries were prepared from two biological replicates from wt male controls, three *Kdm6a*^*Δ/Y*^ and one *Kdm6a*^*ΔP/-Y*^ male ES cell clones as well as one biological replicate from wt female control and one female *Kdm6a*^*ΔP/ΔP*^ clone. For CB derived ES cells libraries were prepared from two biological replicates from wt controls, two *Kdm6a*^*Δ/Y*^ and one *Kdm6a*^*ΔP/ΔP*^ clone. cDNA libraries were constructed for analysis on a NextSeq sequencer and 75bp single-end reads were generated. Diploid gene expression was estimated using Tophat/v2.0.14 (Kim et al., 2013) with default parameters and gene-level expression was normalized using TPM (transcripts per kilobase of exon length per million mapped reads). Differentially expressed genes were determined using DESeq2 analysis (Love et al., 2014) with a false discovery rate (FDR) threshold of 0.05 and a fold-change cutoff of 1.5 was applied to the male ES lines, whereas a p-value threshold of 0.1 and a fold-change cutoff of 1.5 was applied to the female ES lines.

#### Allele-specific expression analysis

To estimate allele-specific gene expression in hybrid cells, a “pseudo-*castaneus*” genome was first assembled by substituting known heterozygous SNPs of *cast* (obtained from Sanger Institute 2015/May, version 5 (Keane et al., 2011) (Table S9) into the BL6 mm10 reference genome. RNA-seq reads were mapped using bowtie/v2.2.5 (Langmead and Salzberg, 2012) to the genome and transcriptome of both the BL6 genome and the pseudo-*cast* genome. Only those reads that mapped uniquely and with a high-quality mapping score (MAPQ ≥ 30) to either the BL6 genome or the pseudo-*cast* genome were kept for further analyses. This resulted in ∼11% and ∼9% of reads mapped uniquely to the BL6 allele and the *cast* allele, respectively (Table S6). As previously described (Berletch et al., 2015), all uniquely mapped and high-quality (MAPQ ≥ 30) reads were segregated into three categories: (1) BL6-SNP reads containing only BL6-specific SNP(s); (2) *cast*-SNP reads containing only *cast*-specific SNP(s); (3) allele-uncertain reads, that is, reads that do not contain valid SNPs (Table S6). For each gene *i*, denote the number of exonic BL6-SNP reads and *cast*-SNP reads be *n*_*i*0_ and *n*_*i*1_, respectively. The BL6-allele expression proportion for gene *i* was then calculated as *p*_*i*0_ = *n*_*i*0_/(*n*_*i*0_ + *n*_*i*1_). To account for the mapping biases between the BL6 and the *cast* alleles, the BL6-allele expression proportion was further adjusted using the average BL6-allele mapping ratio of all autosomal genes *r*_*m*_ = *N*_*A*0_/*N*_*A*1_, where *N*_*A*0_ and *N*_*A*1_ are the number of allele-specific autosomal reads in the BL6 genome and the *cast* genome, respectively. Specifically, the adjusted BL6-allele expression proportion 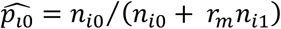. We assume that the diploid TPM value of gene *i* is the sum of haploid TPM values from the BL6-allele and the *cast*-allele. Thus, the BL6-allele TPM is estimated as 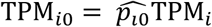, while the *cast*-allele TPM is estimated as 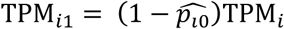. Samples derived from the CB cross were analyzed in a similar manner with the BL6- and pseudo-*cast* reference genomes switched between the maternal and paternal alleles.

Allele-specific differential genes were detected using DESeq2 with similar parameters as described in the above diploid DeSeq2 analysis (Love et al., 2014). Post filtering of allele-specific expression levels was done to remove genes with estimated allelic expression <1 TPM. Allele-specific DEGs were first identified on the maternal-allele and paternal-allele separately, using the correspondingly allelic RNA-seq read counts. Next, allelic down-regulated DEGs were categorized into three groups: genes down-regulated specifically on the maternal allele (Group A), genes down-regulated specifically on the paternal allele (group B), and genes down-regulated from both alleles (Group C), as described in the main text. Similarly, allelic up-regulated DEGs were categorized into three Groups D-F as described in the main text. Note that our allele-specific RNA-seq analyses were performed independently from the diploid RNA-seq analyses, therefore the resulting allelic DEG list does not completely overlap with the diploid DEG list. For example, one gene could be up-regulated from the maternal-allele and down-regulated from the paternal-allele and therefore not identified as a diploid DEG.

#### ATAC-seq with allele-specific analysis

ATAC-seq was done on wt male ES cells and on clone *Kdm6a*^*ΔE1*^ as previously described (Buenrostro et al., 2013). Data were generated from libraries using a NextSeq sequencer to generate 75bp paired-end reads. ATAC-seq reads were mapped to the BL6 genome and the pseudo-*cast* genome, separately, using bwa/ 0.7.12 (Li and Durbin, 2009). Uniquely mapped reads with a high-quality mapping score (MAPQ ≥ 30) to either the BL6 genome or the pseudo-*cast* genome were kept for the following analyses. To assist the allele-specific analysis, ATAC-seq reads were first classified as BL6-SNP reads, *cast*-SNP reads, or allele-uncertain reads, as described in the RNA-seq analysis. As long as one-end of the paired-end read was allele-certain, then the pair was assigned to the same allele. Read pairs with two ends assigned to different alleles were discarded from further analysis. In the following text, we refer to paired-end reads simply as “reads”. Duplicate reads were removed using the MarkDuplicates function in Picard/v2.14.1.

To identify accessible chromatin regions in hybrid mouse cells, diploid ATAC-seq peaks were called by MACS2/v2.1.0 (Zhang et al., 2008) using parameters `--nomodel --shift -50 --extsize 100 --keep-dup all` with p-value cutoff of 0.01. To evaluate DNA accessibility of genes transcribed on the maternal- and paternal-alleles, allelic ATAC-seq read coverage were calculated at promoter regions (2kb upstream and downstream of TSSs). To account for the potential sequencing depth bias, ATAC-seq read coverage were normalized by the number of autosomal reads mapped to the background regions (5kb away from any ATAC diploid peak) on each allele for each library.

#### ChIP-seq with allelic-specific analyses

Cells were harvested and fixed in 1% formaldehyde at room temperature for 10min. Fixation was stopped by adding glycine followed by a 5min incubation at room temperature and cell lysis as described (Nelson et al., 2006). Chromatin was sonicated to yield fragments 300-1000bp in length using a Misonix 3000 sonicator and was then pre-cleared with protein A agarose beads for 1h at 4°C. An aliquot of 20µL was kept to serve as the input fraction. Pre-cleared chromatin was incubated in immunoprecipitation buffer at 4°C overnight using an antibody against H3K27me3. Samples were centrifuged at 1200rpm for 1min and a small portion of the suspension collected as the unbound fraction. Immunoprecipitated chromatin was collected and serially washed in increasingly stringent salt buffers. After elution, crosslinks were reversed in 5M NaCl at 65°C overnight. DNA was purified using Qiaqen mini elute PCR purification kit and subjected to PCR according to the following protocol: 95°C for 3min followed by 35 cycles of 95°C for 30sec, 56°C for 30sec and 72°C 30sec. Samples were incubated at 72°C for 10min and analyzed by gel electrophoresis. ChIP-seq libraries were prepared as described (Bonora et al., 2018).

H3K27me ChIP-seq was performed on BC wt male ES cells and on clone *Kdm6a*^*ΔE1*^ as described (Yang et al., 2015). ChIP libraries were prepared using the TruSeq V2 DNA preparation kit and matching input controls were sequenced by a NextSeq sequencer to produce 75bp paired-end reads. Similar to ATAC-seq analysis (see above), ChIP-seq reads were mapped to the BL6 genome and the pseudo-*cast* genome separately, then uniquely mapped reads with high-quality mapping scores were assigned to the correct allele. Diploid H3K27me3 peaks were identified by MACS2/v2.1.0 (Zhang et al., 2008) using the ChIP sample against the matching input control with parameter setting `--broad --keep-dup all` with p-value < 0.01. Allelic promoter (±2kb of the TSS) coverage were scaled according to the sequencing depth and then normalized by the input control.

## DATA AND SOFTWARE AVAILABILITY

The accession numbers for the RNA-Seq, ChIP-seq and ATAC-seq data reported in this paper.

## Notes

### Competing Interest Statement

The authors have declared no competing interest.

